# *In vitro* antioxidant and anti-inflammatory activities, total phenolic and flavonoid contents of *Anadenanthera colubrina* from Northern Peru

**DOI:** 10.1101/2025.01.24.633472

**Authors:** Edert Abel Samame-Caramutti, Luis Salcedo-Valdez, Silvia Suárez-Cunza

## Abstract

The antioxidant and anti-inflammatory activities of the hydroethanolic extract from bark of *Anadenanthera colubrina* (ACE), as well as its total phenolic (TPC) and flavonoid (TFC) contents, were studied *in vitro*. The ACE showed a high antioxidant activity both in the DPPH^●^ radical scavenging (747.6mg Trolox equivalents/g dry extract) and in the FRAP (435.9mg FeSO_4_ equivalents/g dry extract) chemical assays. The extract (50 to 1000μg/mL) inhibited the lipid peroxidation (formation of TBARS), hemolysis induced by H_2_O_2_, and egg albumin denaturation. No activity was observed in the hypotonicity-induced hemolysis assay. The TPC and TFC were 642.7mg GAE/g and 416.2mg CE/g dry extract, respectively. To the best of our knowledge, these activities were reported for the first time for *A. colubrina* from northern Peru, and suggest that the ACE has notable antioxidant and anti-inflammatory properties, and that the high TPC and TFC could be responsible, at least partially, for these activities.

## Introduction

In Peru, there is a prevalence of chronic inflammatory diseases such as diabetes mellitus, obesity (INEI, 2023), and cancer (CDC Perú, 2023), which require attention since they affect not only the adult population, but also children.

*Anadenanthera colubrina* (Vell.) Brenan (Altschul 1964) is a South American fabaceae that has been widely used in folk medicine because of its anti-inflammatory properties (Júnior et al. 2011) so is considered as a potential source of compounds with biological activity.

One of the most commonly used parts is the bark, mainly in the form of syrup, decoction, infusion and macerated (Monteiro et al., 2006b). Some studies have reported the antioxidant properties of bark (Desmarchelier et al. 1999; Melo et al. 2010; Damascena et al. 2014), which could explain its anti-inflammatory activity both *in vivo* (Santos et al. 2013; Lima et al. 2020) and *in vitro* (Maia et al. 2021, 2022).

*A. colubrina* grows in Peruvian rain and seasonally dry tropical forests between 600 and 2000 m above sea level (Marcelo-Peña et al. 2010). There are two varieties of this species, Cebil (Griseb.) Altschul and Colubrina, and only var. Cebil has been reported in Peru (ILDIS LegumeWeb Database, 2023). Although this species has been extensively used in South-American popular medicine (Albuquerque et al. 2007; Medeiros et al. 2013), in northern Peru it is mainly employed as timber-yielding (Marcelo-Peña et al. 2010). Even though several phytochemical studies have described a number of secondary metabolites with known antioxidant and anti-inflammatory activities in the bark (Santos et al. 2013; Mota et al. 2017), most previous studies are ethnobotanical. Considering that there is scarce scientific information that supports its traditional use (Weber et al. 2011), the mechanisms underlying the biological activity of *A. colubrina* are poorly understood (Delices et al. 2023). Taking into account that edaphic factors modify some plants’ attributes, that the Peruvian soil is varied throughout the territory and that the variety native to Peru has not been previously studied, the aim of this report was to determine for the first time the *in vitro* antioxidant and anti-inflammatory activities of *A. colubrina* bark from northern Peru, as well as its total phenolic (TPC) and flavonoid (TFC) contents.

## Results and discussion

### Physicochemical analysis

The soluble solids content was 1.606 ± 0.3 mg/mL and 1.711 ± 0.2 mg/mL, according to the refractive index and gravimetric analysis, respectively. The pH was 5.5-6.0 (slightly acidic).

### Qualitative phytochemical screening and Total phenolic (TPC) and flavonoid (TFC) contents

As presented in Table S1, the ACE has a high content of phenolic compounds (tannins and flavonoids), low content of glycosides, anthraquinones, anthrones and anthranones, and undetectable or no presence of alkaloids, triterpenes, steroids and saponins.

Some previous studies are not consistent with the results of the phytochemical screening. Silva et al. (2020) reported the presence of alkaloids and saponins, as well as the absence of flavonoids, anthraquinones and glycosides in the bark hydroalcoholic extract of *A. colubrina*. On the other hand, Sá et al. (2016) reported that the bark hydroalcoholic extract has phenolic compounds, tannins and sugars, but flavonoids, steroids and terpenoids were not detected. Additionally, Pessoa et al. (2012) reported the presence of reducing sugars, saponins, triterpenes and steroids. These differences would indicate that edaphic and environmental conditions, as well as the differences in the extraction procedures, have an effect on the biosynthesis or presence of secondary metabolites despite being the same species.

The determination of total phenolics and flavonoids confirmed their abundance, with 642.7 ± 17 mg (3.78 ± 0.1 mmol) gallic acid equivalents (GAE)/g and 416.2 ± 20 mg (1.6 ± 0.08 mmol) catechin equivalents (CE)/g dry extract, respectively; that is, the TPC represents about 65% in GAE of the extract, that include mainly flavonoids (approximately 41% in CE of the extract). The flavonoids/phenolics ratio was 0.65. The results are consistent with previous reports on *A. colubrina* bark, that is, the hydroalcoholic extract is rich in phenolic compounds (Rocha et al. 2017; Maia et al. 2022), which includes both tannins and flavonoids (Mota et al. 2017; Da Costa et al. 2020).

The report of Mota et al. (2017) is particularly relevant since, in addition, they compared the characteristics of the bark of *Anadenanthera colubrina* with those of the other species of the genus, *A. peregrina*. In this respect, such study showed that the hydroethanolic extract of *A. colubrina* has higher TPC (682 mg GAE/g vs. 583 mg GAE/g dry extract, respectively) and TFC (445 mg CE/g vs. 148 mg CE/g dry extract, respectively) than the extract of *A. peregrina*.

Interestingly, the variety of *A. colubrina* native to Northern Peru of the present report possesses properties that are slightly different from those of the Brazilian variety reported by Mota et al. In this regard, the Brazilian variety showed higher TPC and TFC (682 mg GAE/g extract and 445 mg CE/g extract for the Brazilian variety vs. 643 mg GAE/g extract and 416 mg CE/g extract for the Peruvian variety, respectively). In spite of this, as will be seen later, the Peruvian variety has higher antioxidant properties than the Brazilian variety reported by Mota et al.

### Antioxidant activity (Chemical and biological assays)

#### DPPH^●^ free radical scavenging assay

The ACE showed an IC_50_ (4.37 µg/mL) value close to standard Trolox (3.26 µg/mL, Figure S1 and Table S2). The above-mentioned value represents a Trolox equivalent antioxidant capacity (TEAC-DPPH^●^) of 747.55 mg (2.99 mmol) Trolox equivalents/g dry ACE (Table S2).

#### Ferric ion reducing antioxidant power (FRAP) assay

The ACE exhibited an antioxidant capacity of 435.88 mg (2.87 mmol) FeSO_4_ equivalents/g dry ACE (Figure S2 and Table S2).

#### Inhibition of lipid peroxidation using the thiobarbituric acid reactive substances (TBARS) assay

All the concentrations tested (100, 500 and 1000 μg/mL) were effective in reducing the production of TBARS in a dose-dependent manner, by 22, 59 and 68%, respectively, in relation to the induced stress (IS) group (Figure 1a). No difference was found between the ACE treatment at 1000 μg/mL and the Control group (72%).

**Figure 1:**
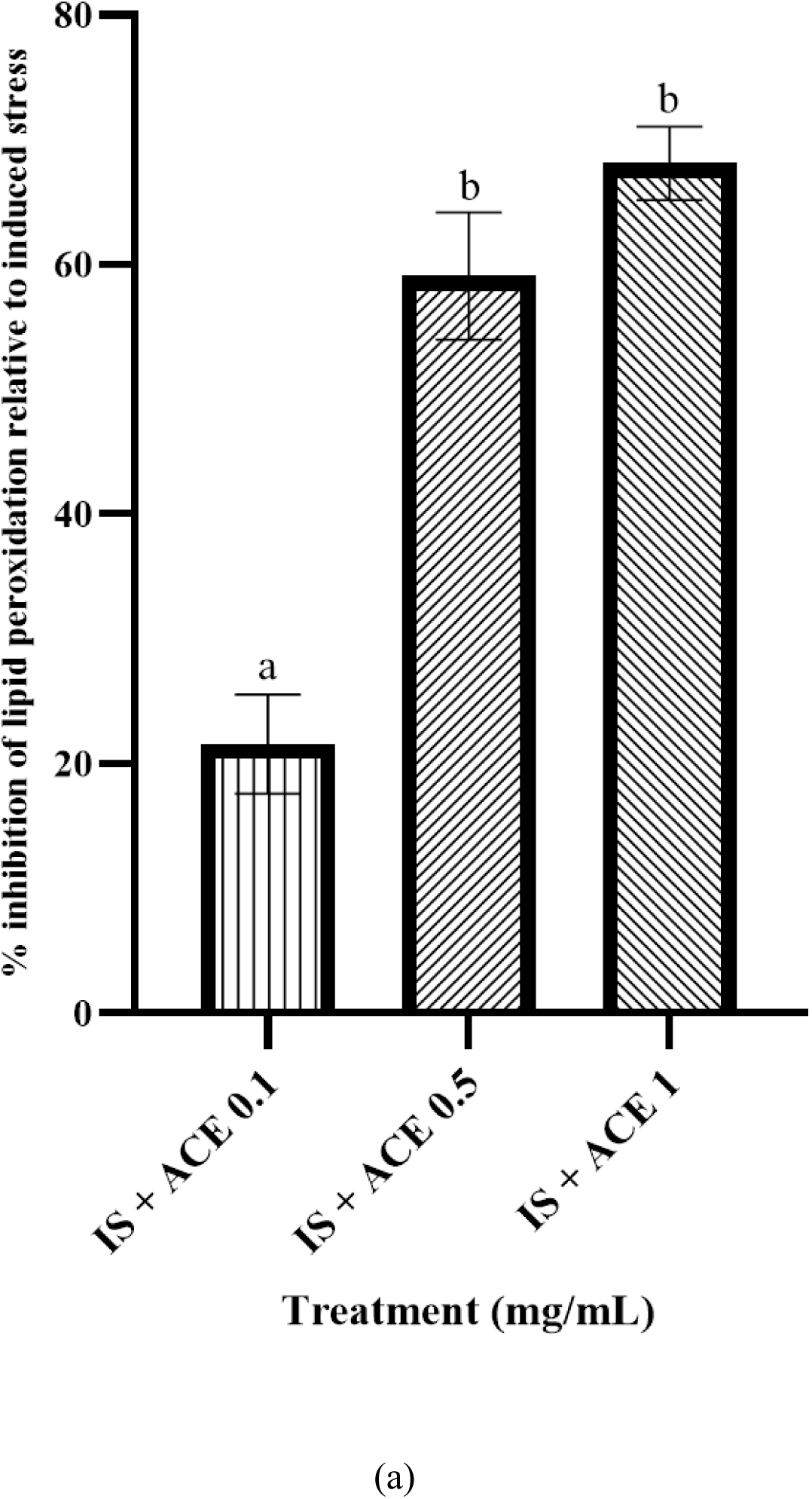
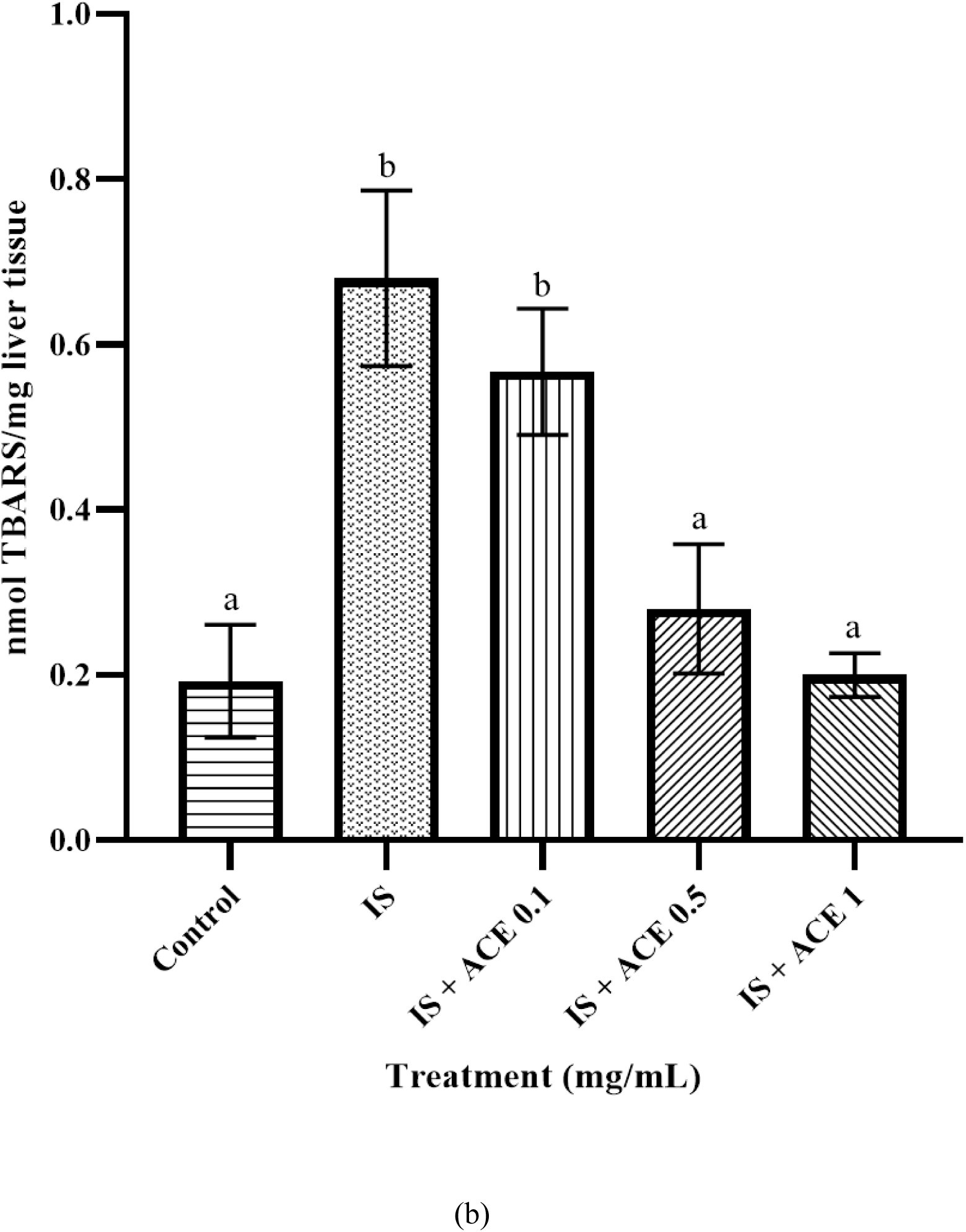
(a) Percentage of inhibition of lipid peroxidation by the hydroethanolic extract from bark of *A. colubrina* (ACE) and (b) effect of the ACE on the formation of TBARS in rat liver homogenate. Results are presented as mean ± SD (*n* = 3), representative of two or three replicates and were evaluated by one-way ANOVA and Tukey test (*p*<0.05). Statistical differences are indicated with different letters. IS: induced stress.

Additionally, the formation of TBARS (in nmol/mg liver tissue) was higher in the IS group than in the ACE treatments at 500 and 1000 μg/mL, indicating a dependence on the concentration of extract. No differences were found neither between the IS group and the ACE at 100 μg/mL, nor between the Control and the ACE treatments at 500 and 1000 μg/mL (Figure 1b).

#### Oxidative hemolysis induced by H_2_O_2_ assay

The ACE, administered at 50, 100 and 150 μg/mL, decreased the hemolysis caused by H_2_O_2_ when compared with the IS group (Figure 2). The prevention of hemolysis depended on the dose only for the 50 and 100 μg/mL treatments (49 and 75% of inhibition, respectively), since the 150 μg/mL treatment caused an inhibition (65%) similar to that observed for the 100 μg/mL group. Interestingly, the percentage of inhibition for the three concentrations of the ACE was much higher than that reported for the Control group (18%).

**Figure 2:**
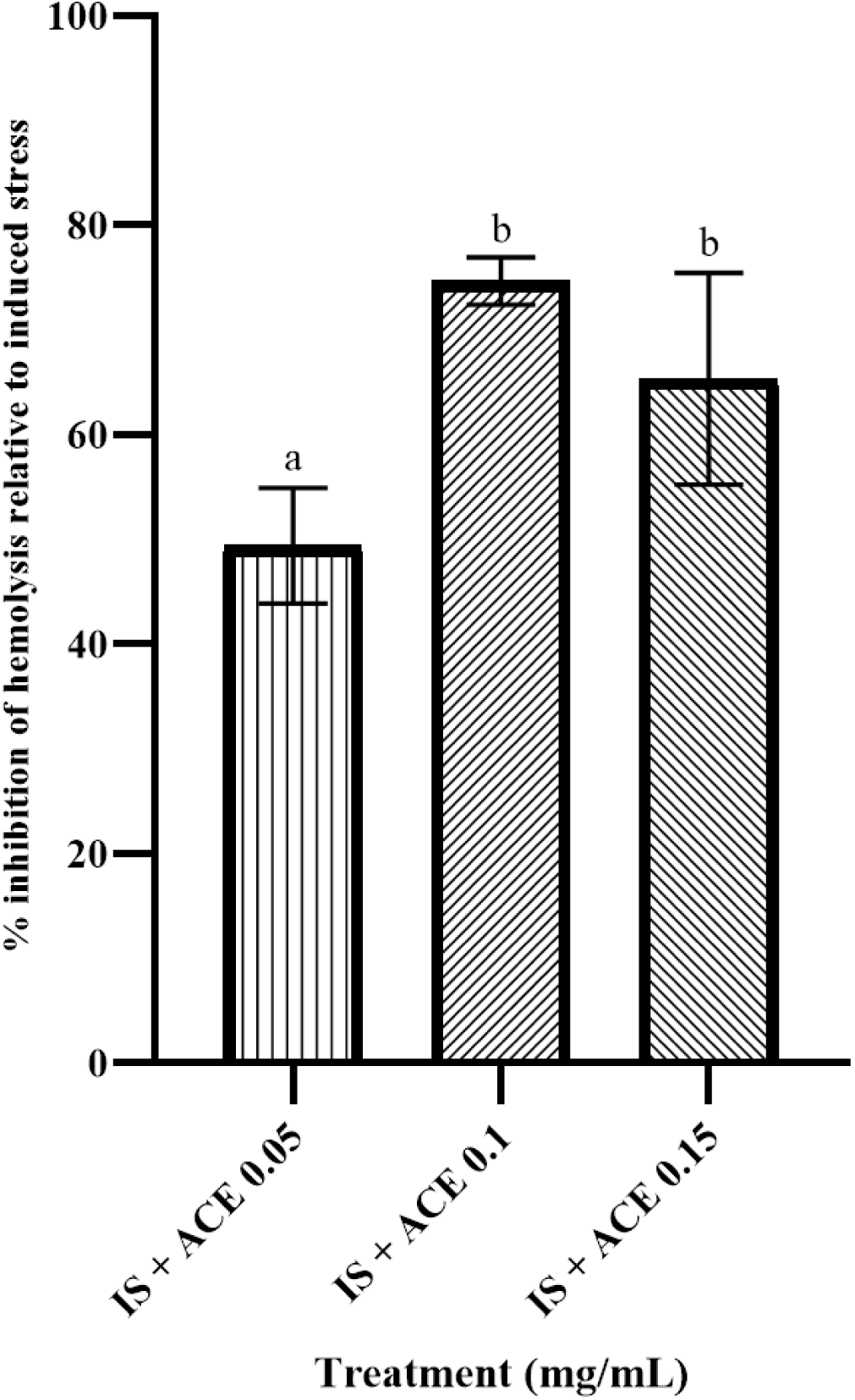
Inhibition of oxidative hemolysis by H_2_O_2_ by the hydroethanolic extract from bark of *A. colubrina* (ACE). Results are presented as mean ± SD (*n* = 3), representative of three replicates and were evaluated by one-way ANOVA and Tukey test (*p*<0.05). Statistical differences are indicated with different letters. IS: induced stress.

Then, the ACE showed a good behavior in two antioxidant assays with different mechanisms of action, single electron transfer for the FRAP assay, and either hydrogen atom or single electron transfer for the DPPH^●^ radical scavenging assay (Prior et al. 2005). Furthermore, as reported by Desmarchelier et al. (1999), the ACE decreased the formation of malondialdehyde and other TBARS as by-products of lipid peroxidation in rat liver homogenates, and additionally reduced the oxidative injury caused by H_2_O_2_ in a human red blood cell (HRBC) model, showing the capacity of the ACE of diminishing the *in vitro* oxidative stress.

These results suggest that the Peruvian variety of *A. colubrina* possesses the antioxidant activity previously reported for the bark (Desmarchelier et al. 1999; Melo et al. 2010; Damascena et al. 2014; Da Costa et al. 2020). The antioxidant capacity against H_2_O_2_ is important, considering that this ROS is a molecule that occurs at sites of inflammation and can move inside the cell and cross membranes, causing cellular damage not only at its site of origin, although the effects of H_2_O_2_ depend on its concentration at the cellular level.

This antioxidant activity is probably due to ACE phytochemical content, mainly phenolic compounds (tannins and flavonoids) (Mota et al. 2017). However, there are some contradictory results regarding the production of phenolics by *A. colubrina* in different environmental conditions (Monteiro et al. 2006a; Araújo et al. 2015), which in turn could explain some discrepancies between the present study and previous reports (Melo et al. 2010; Mota et al. 2017; Silva et al. 2020).

It is interesting that the Peruvian variety exhibited higher TEAC-DPPH^●^ and IC_50_ values than the Brazilian variety reported by the study of Mota et al. (2017) aforementioned (269 mg Trolox equivalents/g extract and 13 μg/mL for the Brazilian variety vs. 748 mg Trolox equivalents/g extract and 4 μg/mL for the Peruvian variety, respectively). Thus, the antioxidant properties (TEAC-DPPH^●^ and IC_50_) of the Brazilian variety were considered only of moderate intensity, whereas those of the Peruvian variety could be considered of high intensity.

These results would confirm that, in addition to the differences in the extraction techniques, edaphic and environmental conditions have an influence over the presence of secondary metabolites and, therefore, on the biological activities of the extracts.

### Anti-inflammatory activity

#### Inhibition of egg albumin denaturation

A dose-dependent inhibition of albumin denaturation was observed for the ACE at 125, 250 and 500 μg/mL by 34, 42 and 47%, respectively, in relation to the IS group (Figure 3). Interestingly, the extract at 750 μg/mL reduced the denaturation only by 40%, lesser than the inhibition observed at 500 μg/mL. In any case, the ACE, at all the concentrations tested, exhibited a higher inhibition than the NSAID diclofenac sodium at 0.5 and 1 mg/mL (19 and 22%, respectively), but lower than the reference at 3 mg/mL (58%). Both the ACE at 250 and 500 μg/mL showed better inhibition of protein denaturation than diclofenac sodium at 2 mg/mL.

**Figure 3:**
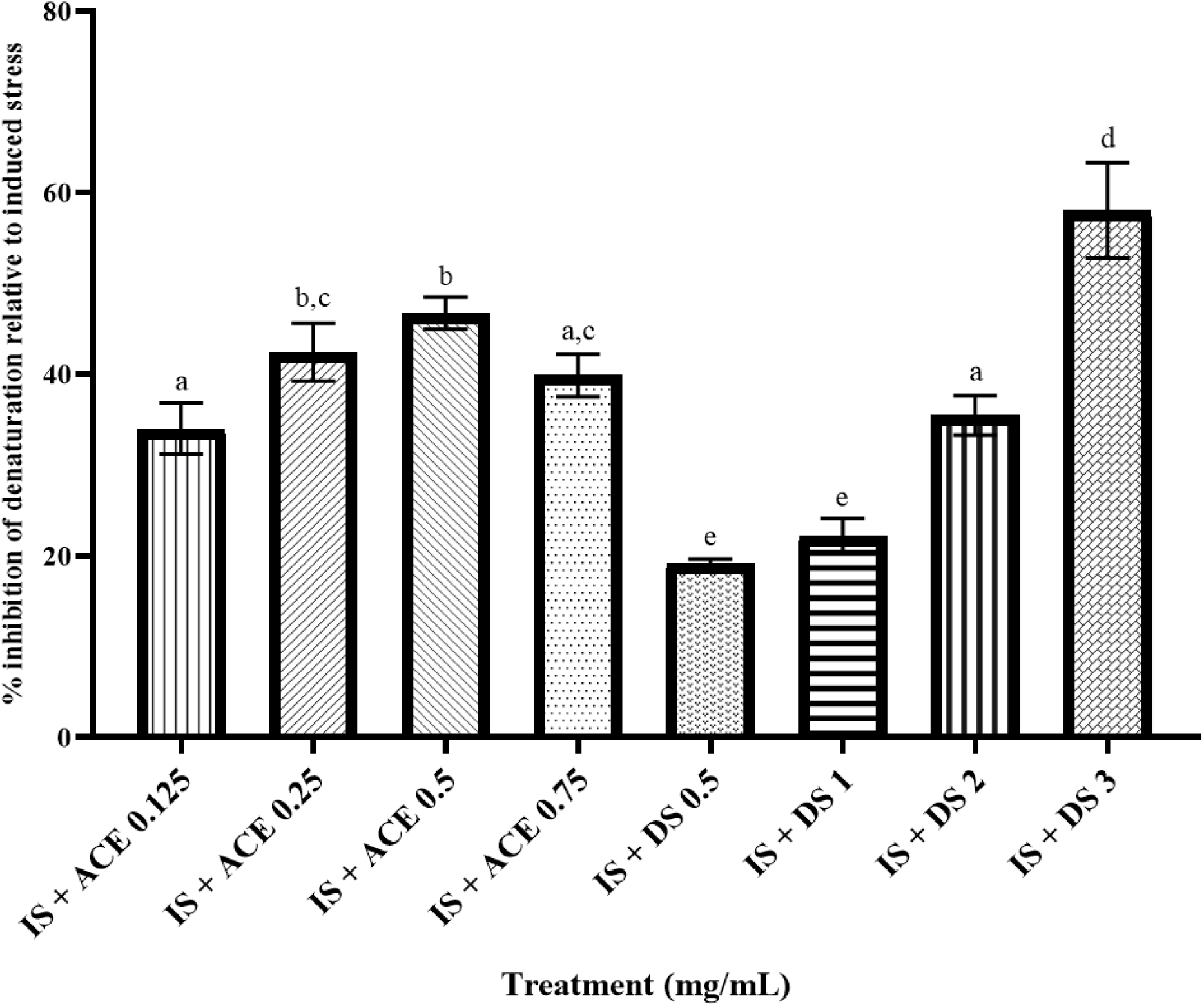
Inhibition of egg albumin denaturation by the hydroethanolic extract from bark of *A. colubrina* (ACE). Diclofenac sodium (DS) was used as standard. Results are presented as mean ± SD (*n* = 3), representative of three replicates and were evaluated by one-way ANOVA and Tukey test (*p*<0.05). Statistical differences are indicated with different letters. IS: induced stress.

#### Stabilization of human red blood cell (HRBC) membrane

The ACE, at 250, 500 and 750 μg/mL, did not show any significant effect on the hemolysis induced by hypotonicity. In this respect, the extract at 250 and 500 μg/mL produced a slightly greater hemolysis than the IS group (6 and 5%, respectively), while the concentration of 750 μg/mL inhibited the hemolysis by 6%. In spite of this, none of the concentrations exhibited any significant difference in relation to the IS group (0% inhibition of hemolysis). On the other hand, diclofenac sodium inhibited the hemolysis between 29 and 42% at the concentrations employed (1, 2 and 3 mg/mL) (Data not shown).

It has been suggested a correlation between the antioxidant and anti-inflammatory activities (Cardoso-Junior et al., 2021). Considering that previous studies have shown that *A. colubrina* bark is rich in metabolites with known anti-inflammatory activity, for example phenolics (Ribeiro et al. 2018), such as gallic acid, catechin, chlorogenic acid and naringin (Da Costa et al., 2020), the anti-inflammatory activity of the Peruvian variety was studied *in vitro*.

The assay of inhibition of egg albumin denaturation is based on the stabilizing action of NSAIDs on the coagulation of proteins subjected to heat (Mizushima and Kobayashi 1968). Given that tissue protein denaturation is able to initiate immune responses (Atassi and Casali 2008) and cause inflammatory disorders, it has been suggested that the compounds that prevent the protein denaturation could have anti-inflammatory potential (Aidoo et al. 2021).

In this regard, the ACE prevented the heat-induced denaturation of egg albumin in a higher degree than the NSAID diclofenac sodium, which have a known protective activity against protein denaturation.

In order to corroborate this anti-inflammatory effect, the stabilization of HRBC membrane assay was performed. This assay is based on the resemblance between the erythrocyte and lysosomal membranes. Considering that lysosomes can release their hydrolytic enzymes into the affected tissues during the inflammatory response, the stabilization of the lysosomal membrane is desirable. The effect of any extract on the stabilization of erythrocyte could be extrapolated to the stabilization of lysosomal membrane and as such, the anti-inflammatory activity of an extract can be evaluated through the prevention of hemolysis (Saleem et al. 2011).

No activity was reported at the same concentrations employed in the denaturation assay. These results are not entirely unexpected, since the ACE is rich in tannins (Table S1), which may have a hemolytic activity on HRBC (Ortega et al. 2006; Deng et al. 2019) as well as erythrocytes from other species (Fedeli et al. 2004). Moreover, the ACE caused hemolysis at concentrations ≥ 250 mg/mL in the H_2_O_2_-induced oxidative stress assay above-mentioned (data not shown). In this respect, Rocha et al. (2017) reported that the hydroalcoholic extract from bark of *A. colubrina* was slightly toxic, even at low concentrations (0.25 to 32 mg/mL), on HRBC.

Although these results would indicate that the ACE is a potential source of anti-inflammatory compounds, especially those related with protein denaturation, it must be considered that the assays were performed *in vitro*, and does not necessarily show what would occur *in vivo*, since bioavailability of active compounds, such as tannins, must be considered. However, it is promising that *A. colubrina* has shown better inhibitory activities than a NSAID such as diclofenac sodium in one of the assays employed.

## Conclusions

To the best of our knowledge, the properties of *Anadenanthera colubrina* were reported for the first time in Peru. The hydroethanolic extract from the bark (ACE) possesses strong *in vitro* antioxidant activity, as well as high contents of total phenolics and flavonoids. Furthermore, the ACE exhibited anti-inflammatory activity in the albumin denaturation assay. These activities could be attributable, at least partially, to its content in bioactive compounds, suggesting that *A. colubrina* from northern Peru is a good source of compounds with biological activity. Further studies are recommended to evaluate the mechanism of action of the ACE.

## Acknowledgements

This work was partially supported by the Universidad Nacional Mayor de San Marcos (UNMSM). The authors thank PhD. Mercedes Soberón-Lozano from the UNMSM and PhD. Consuelo Rojas-Idrogo from the Universidad Nacional Pedro Ruiz Gallo (UNPRG).

## SUPPLEMENTAL SECTION

## Experimental section

### Reagents and standards

2,2-diphenyl-1-picryl-hydrazyl (DPPH), Folin-Ciocalteu’s phenol reagent (2N), gallic acid, (+)-catechin, (±)-6-hydroxy-2,5,7,8-tetramethylchroman-2-carboxylic acid (Trolox), 2-thiobarbituric acid and sodium phosphate monobasic (≥ 99%) were obtained from Sigma-Aldrich (MO, USA). Sodium carbonate (Na_2_CO_3_), sodium chloride (≥ 99.5%), D(+)-glucose, Perhydrol® 30% H_2_O_2_ and disodium hydrogen phosphate anhydrous (99%) were purchased from Merck (Germany). Citric Acid-1-hydrate was obtained from Riedel-de Haën (Germany). Diclofenac sodium was a generous gift from Pharmacist Zoila Gallegos-Salazar from Laboratorios Portugal (Arequipa, Peru). All other chemicals used were of analytical grade.

All the absorbance values were monitored using a Thermo Fisher Scientific Genesys 50S UV-Vis spectrophotometer (Waltham, MA, USA) into 1 cm quartz cuvettes.

### Plant material and extraction

The bark of *Anadenanthera colubrina* (Vell.) Brenan was collected locally in the primary forest of Las Juntas (department of Cajamarca, Peru) (5°20’30” S, 78°46’00” W; mean altitude 625 m) (Tropicos.org, 2024). The plant was authenticated by Prof. Eric Rodríguez-Rodríguez, and two voucher specimens were deposited in the Herbarium Truxillense of the Universidad Nacional de Trujillo (Trujillo, Peru) (voucher code numbers: HUT 57669 and HUT 57670).

The hydroethanolic extract was obtained by methods reported in literature for other fabaceae (Kondeti et al. 2010; Olaleye et al. 2013), with modifications (Samame-Caramutti 2015); briefly, the bark was dried out for 14 days at room temperature and milled. A sample of the powdered bark was extracted by a successive exhaustive extraction with 96% ethanol (1:2 w/v) for 48 hours at room temperature in agitation. The crude extracts were combined, filtered, and concentrated till dryness, obtaining a brownish red dry product (hydroethanolic extract from the bark of *A. colubrina*, ACE). The extraction yield was 20.7%. The ACE was stored at 4°C.

The ACE was completely dissolved in a 96% ethanol: double-distilled water mixture (1:29), and this solution was used for all the assays.

### Physicochemical analysis

The soluble solids content of the ACE (2 mg/mL) was determined gravimetrically, using an Ohaus® Pioneer^TM^ analytical balance (120 g, d = 1 mg; Parsipanny, NJ, USA), and by measuring the refractive index (solvent: 96% ethanol: double-distilled water, 1: 29), using an ATAGO® PAL-α hand refractometer (Tokio, Japan). The pH was obtained using pH-indicator strips (Merck, Germany).

### Qualitative phytochemical screening and Total phenolic (TPC) and flavonoid (TFC) contents

The qualitative phytochemical analysis was performed according to Lock de Ugaz (1994).

The TPC was determined by the Folin-Ciocalteu method (Singleton et al. 1999). The ACE (40-120 µg/mL, 0.1 mL) was incubated with 10% Folin reagent (0.5 mL) and 7.5% Na_2_CO_3_ (0.4 mL) at room temperature for 30 minutes. The changes in absorbance were read at 765 nm. The results were expressed as mg of gallic acid equivalents (GAE)/g of dry ACE, and calculated from a calibration curve using gallic acid as standard (0-100 μg/mL), and the formula:

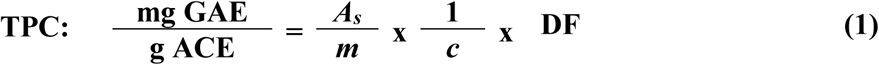

where *A_s_* is the absorbance with the sample (ACE), *m* is the slope from the equation of a straight line of calibration of gallic acid, *c* is the concentration of sample (ACE) and DF is the dilution factor.

The TFC was estimated as described by Zhishen et al. (1999). The ACE (50-250 µg/mL, 0.5 mL) was incubated, in succession, with 5% NaNO_2_ (w/v; 0.15 mL) for 5 minutes, 2.5% AlCl_3_ (w/v; 0.25 mL) for 6 minutes, and 1M NaOH (0.25 mL) for 10 minutes. The changes in absorbance were read at 510 nm. The results were expressed as mg of catechin equivalents (CE)/g of dry ACE, and calculated from a calibration curve using catechin as standard (0-72 µg/mL), and the formula:

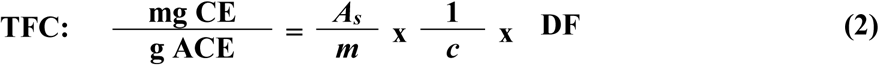

where *m* is the slope from the equation of a straight line of calibration of catechin, and *A_s_*, *c* and DF are as aforementioned.

### Antioxidant activity (Chemical and biological assays)

#### DPPH^●^ free radical scavenging activity

The scavenging activity of DPPH^●^ free radical was evaluated according to Brand-Williams et al. (1995). Briefly, a DPPH^●^ solution (20 mg %) prepared in ethanol was stirred for 40 minutes; this stock solution was kept in dark at 4°C. The initial absorbance of the working solution was adjusted to 0.9 ± 0.02 at 517 nm (DPPH^●^ concentration: 78.5 μM) with the same solvent, and employed to prepare the control (absorbance: 0.6 ± 0.02). The ACE (5-20 µg/mL, 0.4 mL) was incubated with the DPPH^●^ working solution (0.8 mL) for 30 minutes in the dark. The changes in absorbance were read at 517 nm. The percentage of radical scavenging was calculated as

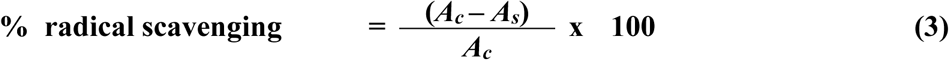

where *A*_c_ and *A*_s_ are the absorbances without and with the sample, respectively.

The values of inhibitory concentration of ACE required to scavenge 50% of the initial free radical (IC_50_) were calculated by linear regression. Trolox equivalent antioxidant capacity (TEAC-DPPH^●^) was expressed as mg TE/g of dry ACE, and calculated from a calibration curve using Trolox as standard (0-15 µg/mL), and the formula:

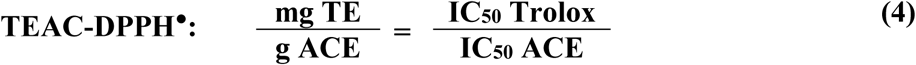

#### Ferric ion reducing antioxidant power (FRAP) assay

The assay was performed according to Benzie and Strain (1996). The ACE (40-160 μg/mL, 50 μL) was added to 950 μL of the FRAP working solution (10mM TPTZ in 40mM HCl, 20mM FeCl_3_ in double-distilled water and 0.3M acetate buffer, pH 3.6, in a ratio of 1:1:98). The reaction mixtures were incubated for 10 min. The changes in absorbance were read at 593 nm.

The results were expressed as mg FeSO_4_ equivalents/g of dry ACE, and calculated from a calibration curve using FeSO_4_ as standard (0-91 µg/mL), and the formula:

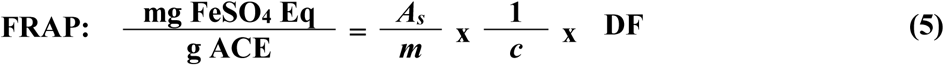

where *A_s_* is the absorbance with the sample (ACE), *m* is the slope from the equation of a straight line of calibration of FeSO_4_, *c* is the concentration of sample (ACE) and DF is the dilution factor.

#### Inhibition of lipid peroxidation using the thiobarbituric acid reactive substances (TBARS) assay

The oxidative stress, expressed as TBARS, was measured by the method of Buege and Aust (1978), with modifications. A male Hotzmann rat (200-250 g), previously anesthetised, was sacrificed by cervical dislocation. The liver was removed, perfused with cold 0.154M KCl, and homogenized in cold phosphate-buffered saline (PBS, pH 7.4). The lipid peroxidation was initiated by adding to 5% liver homogenate (w/v, 0.9 mL), successively, 4mM ascorbic acid (0.045 mL) and 2mM FeSO_4_ (0.015 mL). Afterwards, different concentrations of either ACE (100-1000 μg/mL, 0.06 mL) or PBS (pH 7.4, 0.06 mL and 0.12 mL for the induced stress and control groups, respectively) were included in the reaction mixtures; then, these were incubated for 30 minutes at room temperature. Subsequently, each reaction mixture (0.3 mL) was mixed with 20% trichloroacetic acid (0.6 mL), heated in boiling water for 15 minutes, cooled down with tap water, mixed with 0.67% thiobarbituric acid (TBA in 0.25N HCl, 0.9 mL) and heated again in boiling water for 30 minutes. After cooling down with cold tap water, each tube was centrifuged (5,488xg for 10 minutes), the supernatant was collected and the absorbance was determined at 535 nm.

The antioxidant activity was expressed as percentage of inhibition of lipid peroxidation and as concentration of TBARS:

The percentage of inhibition of lipid peroxidation was calculated as follows:

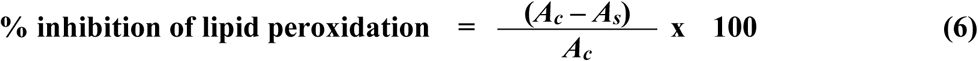

where *A_c_* is the absorbance of the induced stress group (0% inhibition of lipid peroxidation); and *A_s_* is the absorbance with the sample (ACE).

The concentration of TBARS was expressed as nmol TBARS/mg liver tissue, and was calculated as follows:

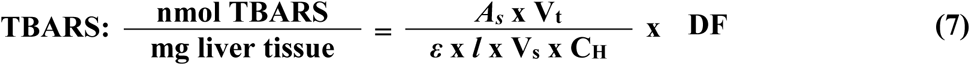

where *A_s_* is the absorbance of each reaction mixture, V_t_ is the final volume of the reaction mixture, *ε* is the molar absorption coefficient of the MDA-TBA_2_ complex at 535 nm (1.56 x 10^5^ M^-1^ cm^-1^), *l* is the path length of the cuvette (1 cm), V_s_ is the volume of the sample in the reaction mixture, C_H_ is the concentration of rat liver homogenate and DF is the dilution factor.

#### Oxidative hemolysis induced by H_2_O_2_ assay

The assay was performed following the method of Xu et al. (2014), with modifications. Fresh human venous blood (3 mL) was collected from healthy donors and diluted with sterilized Alsever’s solution (2% dextrose, 0.8% sodium citrate, 0.05% citric acid and 0.42% NaCl in double-distilled water). The solution was centrifuged (625xg for 3 minutes) and washed three times with the same volume of solvent (cold 1X PBS, pH 7.4) to separate the human red blood cells (HRBC) from the white cell layer, plasma, and free hemoglobin (Hb) released from the injured HRBC. A 5% HRBC suspension (in cold Alsever’s solution, 0.1 mL) was mixed with either ACE (50-150 μg/mL, 0.45 mL) or solvent (0.45 mL and 0.9 mL for the induced stress and control groups, respectively). The reaction mixtures were pre-incubated at 37°C for 15 minutes. Then, 166mM H_2_O_2_ (in solvent, 0.45 mL) was added, the reaction mixture was incubated at 37°C for 4 hours, and centrifuged at 625xg for 10 minutes. The Hb content in the supernatant was estimated at 405 nm. After every trial, the HRBC solution was stored at 4°C for not more than 72 h to ensure cell viability and cellular antioxidant level.

The percentage of prevention of hemolysis was calculated as follows:

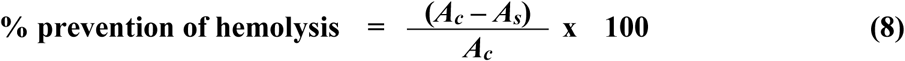

where *A*_c_ is the absorbance of the induced stress group (0% prevention of hemolysis), and *A*_s_ is the absorbance with the sample (ACE).

### Anti-inflammatory activity

#### Inhibition of egg albumin denaturation

The assay was carried out as described by Mizushima and Kobayashi (1968) and Chandra et al. (2012), with modifications. The reaction mixture (1.3 mL) consisted of fresh egg albumin: PBS (pH 6.4) (1:1, 0.1 mL), PBS (pH 6.4, 0.7 mL), and different concentrations of the ACE (125-750 μg/mL, 0.5 mL). Double-distilled water and diclofenac sodium (0.5-3 mg/mL) were used for the induced stress group and as standard, respectively. The reaction mixtures were incubated at 37 ±1°C for 20 minutes, and then heated at 65 ± 2°C for 5 minutes to induce denaturation. After cooling down, the changes in absorbance were read at 660 nm. The percentage of inhibition of protein denaturation was calculated as

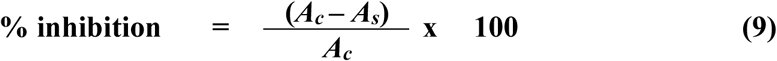

where *A*_c_ is the absorbance of the induced stress group (0% inhibition of protein denaturation), and *A*_s_ is the absorbance with the sample.

#### Stabilization of human red blood cell (HRBC) membrane

The assay was performed according to the procedures described by Lavanya et al. (2010) and Torres Carro et al. (2016), with modifications. Fresh human venous blood (3 mL) was obtained from healthy donors, and diluted with an equal volume of sterilized Alsever’s solution. The solution was centrifuged (5,488xg for 10 minutes), and the packed cells were washed five times with an equal volume of isotonic saline (0.85% NaCl, pH 7.2) in order to prepare a 10% (v/v) HRBC suspension with isotonic saline. The suspension was stored at 4°C. The reaction mixture (1.5 mL) consisted of the 10% HRBC suspension (0.2 mL), 0.14M sodium phosphate buffer (pH 7.4, 0.333 mL), and different concentrations of the ACE (250-750 μg/mL, 0.267 mL). Double-distilled water (0.7 mL) was used to induce hypotonic hemolysis. Solvent (96% ethanol: double-distilled water, 1: 29) and diclofenac sodium (1-3 mg/mL) were used for the induced stress group and as standard, respectively. The reaction mixtures were incubated at 37 ± 1°C for 30 minutes, and then centrifuged at 650xg for 3 minutes. The Hb content in the supernatant was determined at 550 nm. After every trial, the HRBC solution was stored at 4°C for not more than 72 h to ensure cell viability.

The percentage of prevention of hemolysis was calculated as follows:

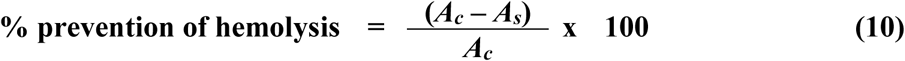

where *A*_c_ is the absorbance of the induced stress group (0% prevention of hemolysis), and *A*_s_ is the absorbance with the sample.

### Statistical analysis

The assays were performed in three independent experiments in duplicate or triplicate. The data were presented as means ± SD. To determine the statistical differences, the data were evaluated with one-way ANOVA followed by the Tukey test. All analyses were made using the Minitab Statistical Software and the GraphPad Prism software (version 8.0.1). The differences were considered significant at *p* < 0.05.

---

**Table S1:** Qualitative phytochemical screening of the hydroethanolic extract from bark of *Anadenanthera colubrina* (ACE)

| Secondary metabolite | Reactive | Result |
| --- | --- | --- |
| Flavonoids | Shinoda | (+++) |
| Phenols | FeCl<sub>3</sub> | (+++) |
| Tannins | Gelatin | (+++) |
| Alkaloids | Dragendorff | (-) |
|  | Mayer | (-) |
|  | Bertrand | (-) |
| Glucosides | Vanillin – sulfuric acid | (+) |
| Anthraquinones, anthrones and anthranones | Bornträger | (+) |
| Triterpenes and/or steroids | Liebermann – Burchard | (-) |
| Saponins | Afrosimetric index | (-) |

Key: (+): Present; (++) : Rich; (+++): Very rich; (-): Undetectable or absent

**Table S2:**
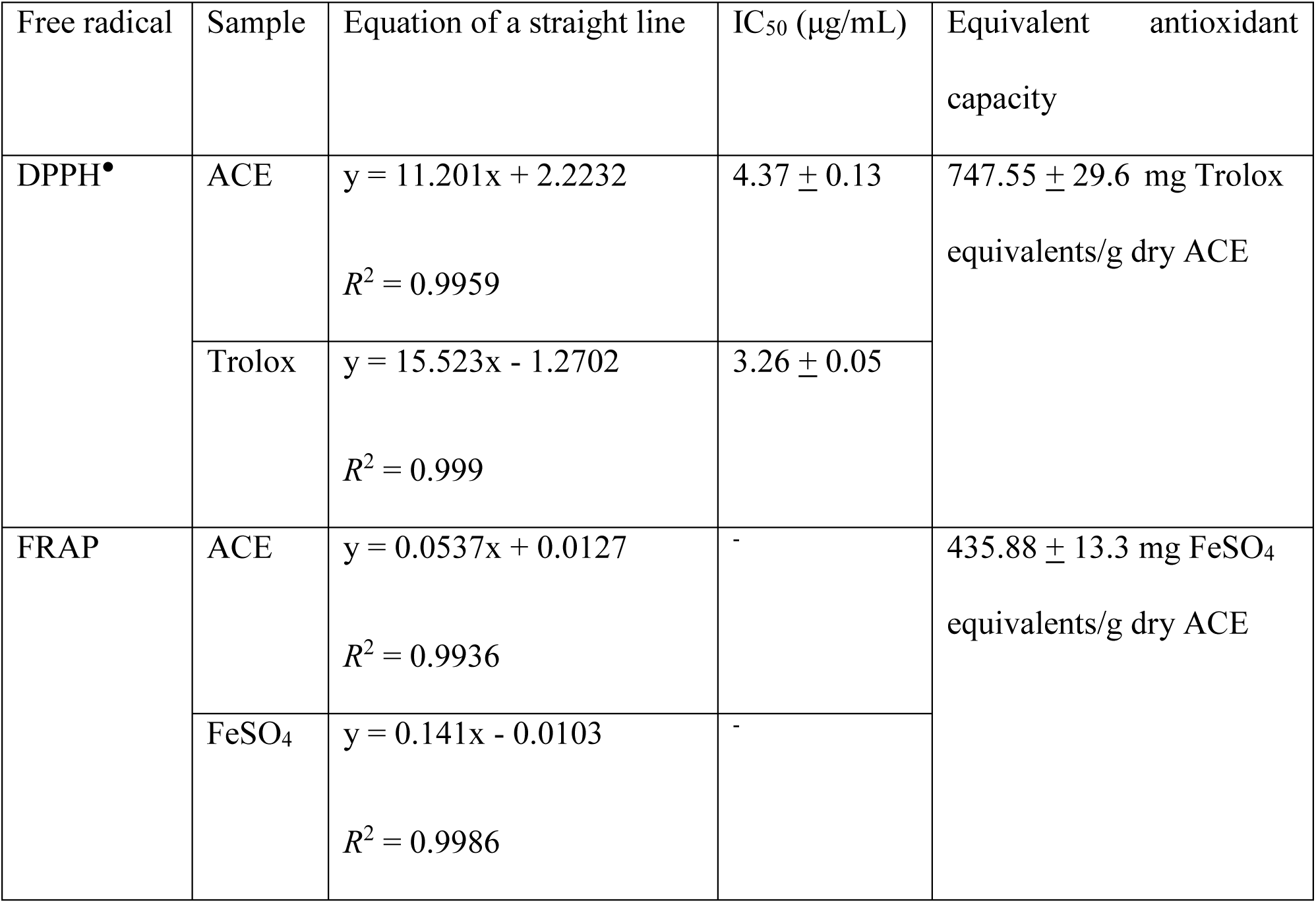
DPPH^●^ radical scavenging activity and ferric ion reducing antioxidant power (FRAP) of the hydroethanolic extract from bark of *A. colubrina* (ACE) and standards. Results presented as mean ± SD (*n* = 3), representative of two replicates.

| Free radical | Sample | Equation of a straight line | IC<sub>50</sub> (μg/mL) | Equivalent antioxidant capacity |
| --- | --- | --- | --- | --- |
| DPPH• | ACE | $y = 11.201x + 2.2232$<br>$R^2 = 0.9959$ | $4.37 \pm 0.13$ | $747.55 \pm 29.6$ mg Trolox equivalents/g dry ACE |
| | Trolox | $y = 15.523x - 1.2702$<br>$R^2 = 0.999$ | $3.26 \pm 0.05$ | |
| FRAP | ACE | $y = 0.0537x + 0.0127$<br>$R^2 = 0.9936$ | - | $435.88 \pm 13.3$ mg FeSO<sub>4</sub> equivalents/g dry ACE |
| | FeSO<sub>4</sub> | $y = 0.141x - 0.0103$<br>$R^2 = 0.9986$ | - | |

**Figure S1:**
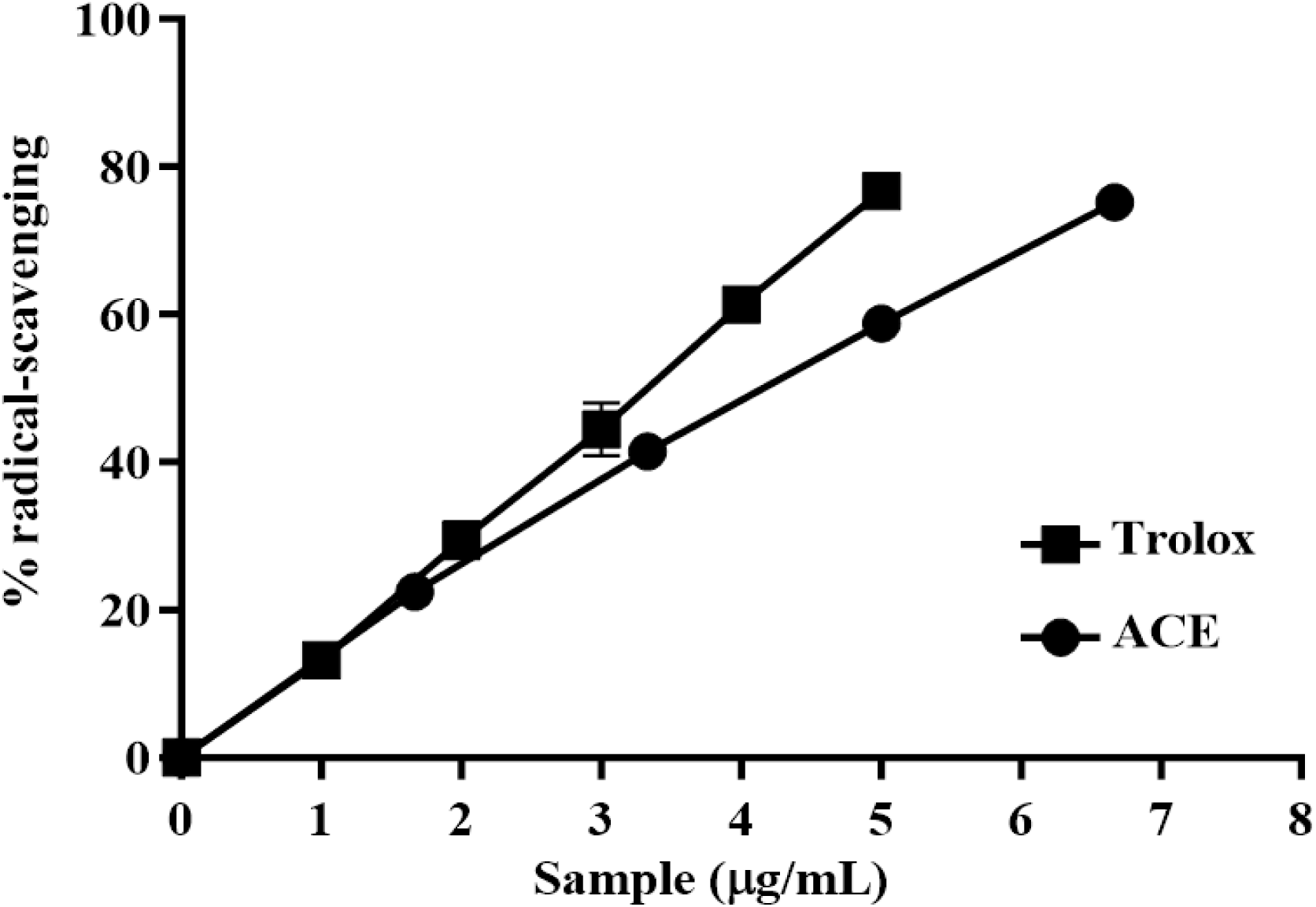
DPPH^●^ (78.5 μM) radical scavenging effects of the hydroethanolic extract from bark of *A. colubrina* (ACE) and Trolox. Results are presented as mean ± SD (*n* = 3), representative of two replicates.

**Figure S2:**
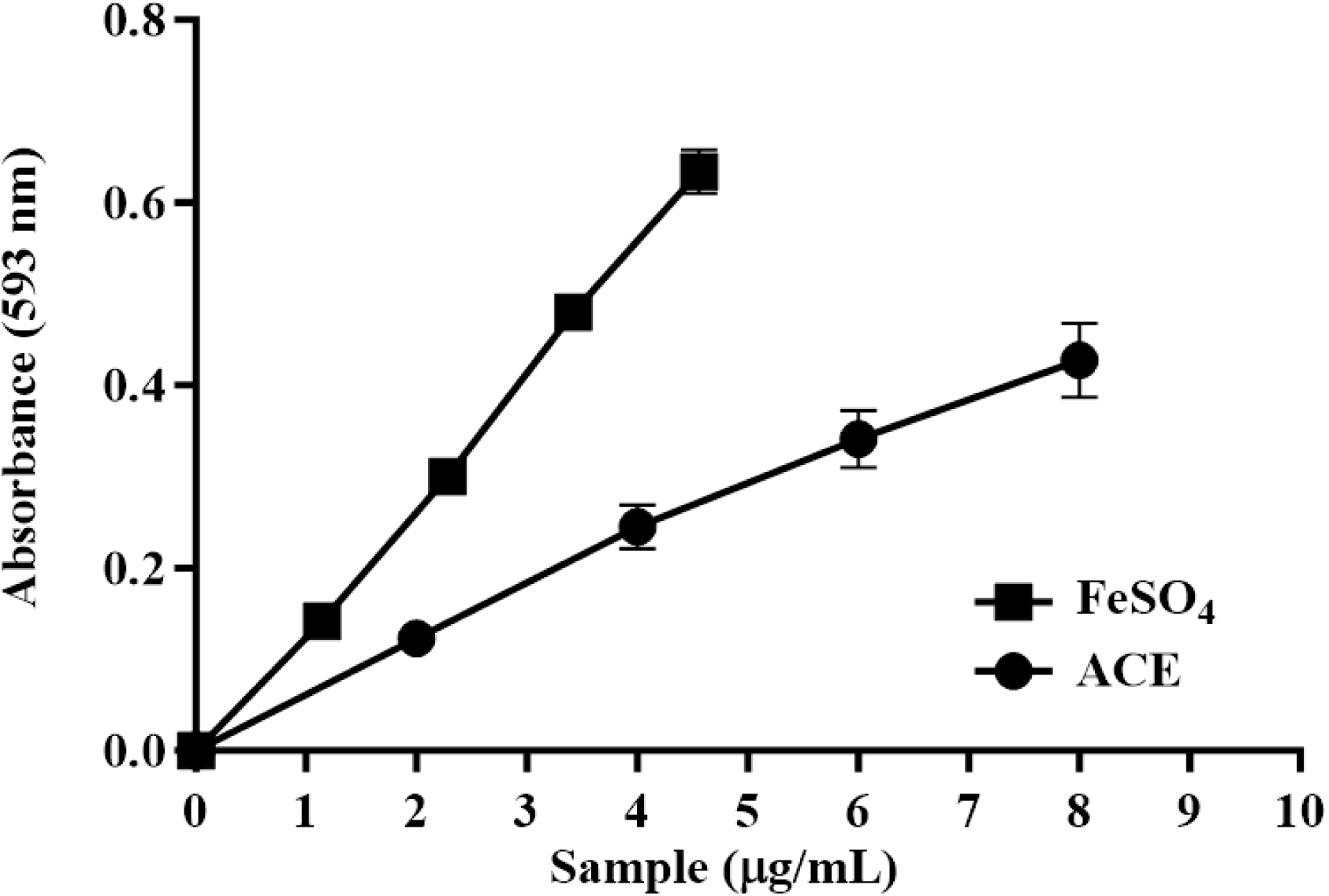
Reactivity of the hydroethanolic extract from bark of *A. colubrina* (ACE) and ferrous sulfate in the FRAP assay. Results are presented as mean ± SD (*n* = 3), representative of two replicates.

